# Cortical networks relating to arousal are differentially coupled to neural activity and hemodynamics

**DOI:** 10.1101/2022.12.01.518759

**Authors:** Lisa Meyer-Baese, Arthur Morrissette, Yunmiao Wang, Brune Le Chatelier, Peter Borden, Shella Keilholz, Garrett Stanley, Dieter Jaeger

## Abstract

Even in the absence of specific sensory input or a behavioral task, the brain produces structured patterns of activity. This organized activity is modulated by changes in arousal. Here, we use wide-field voltage imaging to establish how arousal relates to cortical network voltage and hemodynamic activity in spontaneously behaving head-fixed male and female mice expressing the voltage-sensitive fluorescent FRET sensor Butterfly 1.2. We find that global voltage and hemodynamic signals are both positively correlated with changes in arousal with a maximum correlation of 0.5 and 0.25 respectively at a time lag of 0 seconds. We next show that arousal influences distinct cortical regions for both voltage and hemodynamic signals. These include a broad positive correlation across most sensory-motor cortices extending posteriorly to the primary visual cortex observed in both signals. In contrast, activity in prefrontal cortex is positively correlated to changes in arousal for the voltage signal while it is a slight net negative correlation observed in the hemodynamic signal. Additionally, we show that coherence between voltage and hemodynamic signals relative to arousal is strongest for slow frequencies below 0.15 Hz and is near zero for frequencies greater than 1Hz. We finally show that coupling patterns are dependent on the behavioral state of the animal with correlations being driven by periods of increased orofacial movement. Our results indicate that while hemodynamic signals show strong relations to behavior and arousal, these relations are distinct from those observed by voltage activity.

**Significance Statement:** We leverage wide-field voltage imaging to examine the relation between cortical changes in membrane potential dynamics and hemodynamics. These two signals are then examined with respect to changes in arousal, as measured by pupil diameter, in awake head fixed mice. Our results show similarities as well as important differences in the correlation of arousal with neuronal population activity dynamics and the hemodynamic signal. Further, the spatial activity correlation maps with arousal depended differentially on the behavioral state of the animal in a frequency dependent manner. Our results indicate that the modulation of brain networks by arousal is dynamically regulated, and only partly overlap between functional networks determined from hemodynamic or voltage activity.

## Introduction

The cerebral cortex shows constant activity even in the absence of overt behaviors. This spontaneous resting state activity has been used in humans, primates, and rodents to study the functional architecture of the brain (Biswal, Zerrin Yetkin, Haughton, & Hyde, 1995; Fox, Zhang, Snyder, & Raichle, 2008; Meyer-Baese, Watters, & Keilholz, 2022). Distinct synchronization existing between brain regions is interpreted to represent the connectivity of intrinsic functional neural networks. The study of these resting state functional connectivity networks is predominantly done using functional magnetic resonance imaging (fMRI). Despite its widespread use, fMRI is dependent on measured changes in the blood oxygen level-dependent (BOLD) signal which serves as a surrogate measurement of neural activity (Buxton, 2013). A fundamental assumption in this case is that spontaneous hemodynamic signals are related to neural activity through a neurovascular coupling mechanism that is similar across various cortical areas and behavioral states (Gao et al., 2017).

Behavioral states such as changes in movement and pupil diameter are proposed to be encoded in spontaneous resting state fluctuations (McGinley et al., 2015; Reimer et al., 2014). Previous studies have validated pupil diameter changes as a measure for arousal both in human and animal models (Alnaes et al., 2014; DiNuzzo et al., 2019; P. R. Murphy, O’Connell, O’Sullivan, Robertson, & Balsters, 2014; Reimer et al., 2016). In rodents, a direct relationship is known to exist between increases in arousal and increases in pupil diameter at a short time lag (McGinley et al., 2015; Reimer et al., 2014; Shimaoka, Harris, & Carandini, 2018; Vinck, Batista-Brito, Knoblich, & Cardin, 2015). Animal movement, often measured by periods of locomotion, has also been observed to influence arousal levels. Such locomotion leads to long dilations in pupil diameter and periods of desynchronized cortical activity (McGinley M.; Shimaoka et al., 2018; Vinck et al., 2015). Quiescent periods that exist between bouts of exploratory behaviors are also known to represent transitions in cortical activity, with these state transitions tightly linked to changes in pupil diameter (Reimer et al., 2014). Recent findings suggest that there are observable differences in neurovascular coupling as a function of the animal’s behavioral state (Winder, Echagarruga, Zhang, & Drew, 2017) making it critical to determine how behavioral state is represented spatially across multiple cortical areas for both neural activity and hemodynamics.

In this work, we examined the detailed relationship between neuronal population activity and hemodynamics with respect to changes in pupil diameter and spontaneous orofacial movements in waking mice. We used wide-field voltage imaging in voltage-sensitive fluorescent protein (VSFP) expressing mice using the Butterfly 1.2 sensor (W. Akemann et al., 2012). Butterfly 1.2 mice were crossed with EMX1 cre to yield strong expression across the dorsal cortex. Butterfly 1.2 imaging reflects changes in subthreshold population-level activity in excitatory neurons biased towards the upper layers of dorsal cortex (Walther Akemann et al., 2012) while the hemodynamic signal reflects the change in total hemoglobin from the superficial blood vessels (Ma, Shaik, et al., 2016). We simultaneously monitored changes in pupil diameter and orofacial movements to enable us to parse their spatial and temporal contributions on cortical activity.

Our results show similarities as well as important differences in the correlation of arousal with neuronal population voltage activity and the hemodynamic signal. We find that global signal changes for voltage and hemodynamic activity are both positively correlated with changes in pupil diameter. We next show that arousal influences distinct cortical regions for both voltage and hemodynamic signals. These two signals are both positively correlated with pupil diameter across most sensory-motor cortices extending posteriorly to the primary visual cortex observed in both signals. In contrast, activity in prefrontal cortex is positively correlated to changes in pupil diameter for the voltage signal while it is a slight net negative correlation observed in the hemodynamic signal. Our results indicate that arousal-related brain networks are dynamically regulated, and only partly overlap between functional networks determined from hemodynamic or voltage activity.

## Materials and Methods

In this study, we used a voltage-sensitive fluorescent protein to record both membrane voltage and hemodynamic activity bilaterally across a large extent of mouse dorsal cortex. Unlike traditional voltage-sensitive dye imaging, voltage-sensitive fluorescent proteins enable us to record from cell-type specific circuits (W. Akemann et al., 2012; Isabelle Ferezou, Bolea, & Petersen, 2006). The voltage-sensitive fluorescent protein used here is optimized to show the greatest changes in fluorescence in the sub-threshold voltage range (W. Akemann et al., 2012) which is important when studying the influence of pupil diameter on cortical activity because these changes may be primarily coupled across sub-threshold fluctuations as observed in intracellular recordings (McGinley M., 2015; McGinley et al., 2015). The hemodynamic signal captured here is most sensitive to changes in total hemoglobin which is similar but not identical to BOLD, which is sensitive to changes to deoxyhemoglobin concentration (Ma, Shaik Mohammed, et al., 2016).

### Animal model

Mice expressing the VSFP-Butterfly 1.2 voltage indicator in excitatory neurons in all layers of cortex were bred at Emory University. To do so, Camk2a-tTA;Ai78(TITL-VSFPB) double-cross Tg mice were obtained from Dr. Hongkui Zeng. These mice were further crossed with a homozygous Emx1-cre line expressing cre in excitatory neurons across all cortical layers (Jax 005628). Triple positive mice robustly express VSFP Butterfly 1.2 fluorescence throughout the cell bodies and neuropil of excitatory neurons in the entire cerebral cortex (Figure 1A). Note that VSFP mice can also be derived from first crossing the cre/tTa dependent VSFP-Butterfly reporter line (Madisen et al., 2015) (Ai78-Jax 023528) with a CamK2a-tTa line expressing tTa in excitatory neurons (Jax 007004), followed by a cross with the EMX1 cre-line.

**Figure 1.**
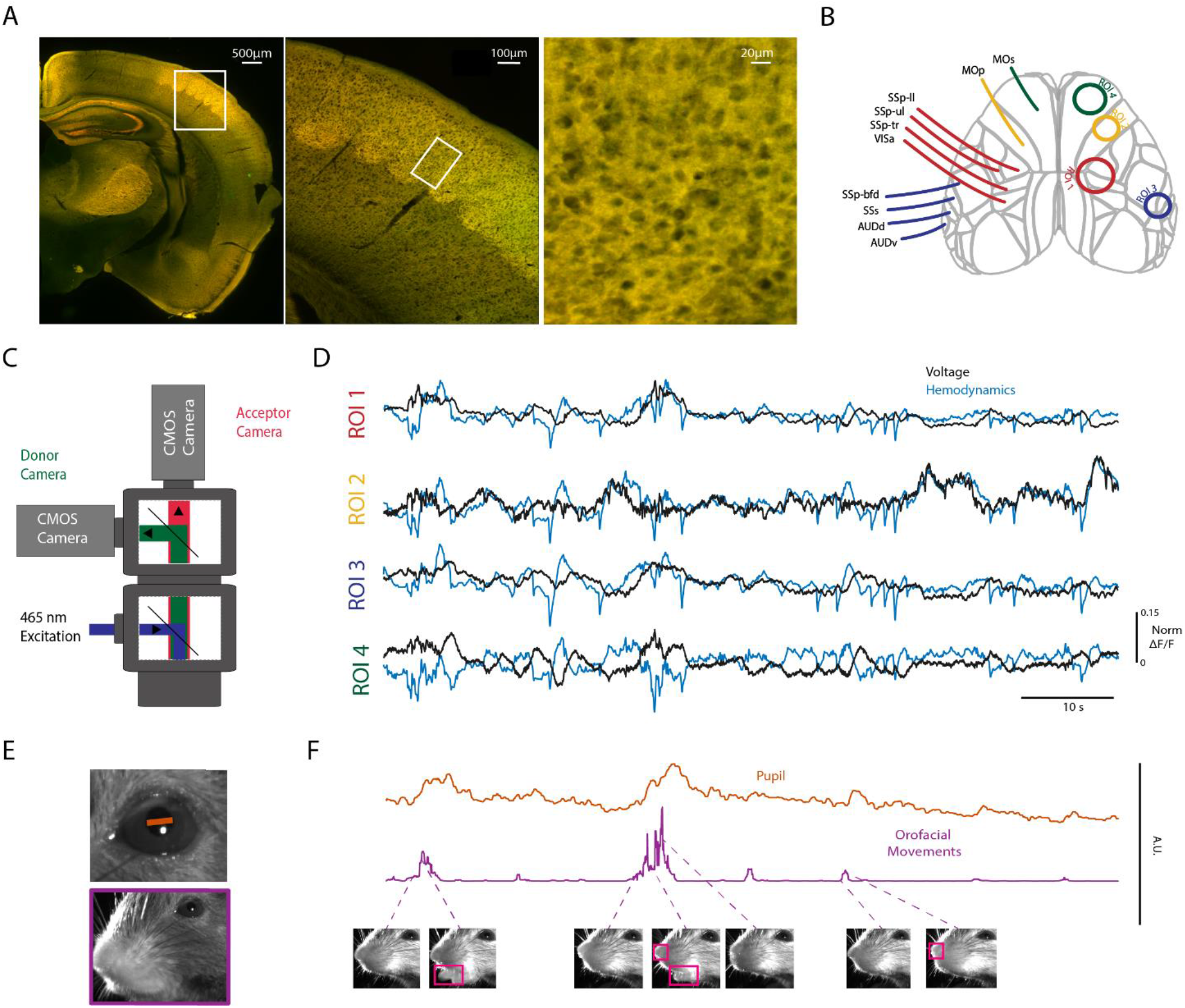
Expression of VSFP protein along with imaging set-up and raw data. **A.** A coronal section of one hemisphere at bregma -1.5 mm depicting combined red and green fluorescence channels reveals cortical VSFP Butterfly 1.2 expression across dorsal cortex. Barrel cortex (white box in left panel) is expanded in resolution to show the membrane associated expression of Butterfly 1.2. Note that voltage imaging is not dependent on even expression of VSFP 1.2 across areas since local ratiometric signals are evaluated. **B.** Cortical map derived from the Allen Brain Mouse Atlas v3. Four ROIs that contained regional differences, used in subsequent analysis. **C.** Imaging set-up to record voltage activity across wide extents of dorsal cortex, using an epi-illumination macroscope containing a 482nm excitation LED and two cameras to capture fluorescent light for both green and red wavelengths, which corresponds to the donor and acceptor fluorophores of the FRET based VSFP-Butterfly 1.2 sensor. **D.** Example traces from one trial showing the average voltage and hemodynamic activity for 4 ROIs depicted in B. **E.** The top image shows the cropped region of interest for calculating the pupil diameter during cortical imaging. The red line represents the maximum width across the pupil used as the measure of pupil diameter. The bottom image shows an example frame from video acquisition of the mouse face during cortical imaging. The region of interest used for analysis of spontaneous facial movements is depicted by the surrounding purple box. Visible areas of the mouse face contributing to movement signal include the nose, snout, jaw, and whiskers. **F.** Example traces of the pupil diameter and orofacial movement from the same trial depicted in D. Frames at the bottom show different types of orofacial movement that the analysis is sensitive to. Pink boxes highlight changes which includes in this example movement of the tongue/jaw, twitches of the nose and a combination of both. For all panels that contain the hemodynamic signal, it has been inverted and lowpass filtered at 5 Hz.

### Voltage imaging

For voltage imaging of excitatory neurons, mice (n=5, 3 male and 2 female) were prepared with a clear-skull window as previously described (Silasi, Xiao, Vanni, Chen, & Murphy, 2016). The cranial window consisted of a custom-cut glass coverslip cemented over the cleaned skull surface. A thin layer of cyanoacrylate glue was placed in between the bone and cement interface. Mice were administered 0.1mg/kg of buprenorphine SR as a long-lasting analgesic before starting the surgery and anesthetized with 1-2% isoflurane (3-4% for induction). All experimental procedures were approved by the Emory University Institutional Animal Care and Use Committee.

VSFP signals were imaged from awake, head-fixed mice, using a macroscope (MiCAM Ultima, Brainvision Inc.) based on the tandem lens and epi-illumination system design (Ratzlaff & Grinvald, 1991). The imaging plane of the macroscope was focused through the intact skull slightly below (∼100μm) the most superficial blood vessels. Excitation light was provided by a blue LED (LEX2-B, Brainvision Inc.), through a band-pass filter at 482nm (FF01-482/35, Semrock Inc.) and a dichroic mirror (FF506-Di03, Semrock Inc.). VSFP-Butterfly FRET is a unimolecular sensor that contains two different-colored chromophores which act as the FRET donor (mCitirine) and acceptor (mKate2). The emission of mCitirine and mKate2 was imaged through two CMOS cameras (MiCAM-Ultima). The first camera recorded the emitted green fluorescence from mCitrine, which was reflected by a second dichroic mirror (FF580-Di03, Semrock, Inc.) and passed through an emission filter (FF01-542/27, Semrock Inc.). The second camera recorded the emitted red fluorescence from mKate2, passed through the second dichroic mirror and an emission filter (BLP01-594R-50, Semrock Inc.). The camera acquisition was controlled by the MiCAM Acquisition Software. Acquisition frame rates varied between 25-200Hz with a spatial resolution of 100×100μm per pixel for a 100×100 pixel array. The power of the LED light was 0.059 mW/mm^2^.

An additional camera was set-up to monitor and record changes in behavioral state and pupil diameter. This was done by acquiring a 10-25Hz video of the side of the mouse face during each imaging session. The camera was positioned to capture the left side of the mouse’s face as well as the left eye and the area where the forelimbs often rest underneath the mouse’s body.

Animals were put on water restriction and habituated to the head fixation set-up gradually over the course of 4 days (Guo et al., 2014). All 5 mice were imaged during multiple imaging sessions for multiple trials (n = 65 trials in total, mean per mouse = 13, std = 4) where each imaging trial ranged from 20 to 160 seconds (mean = 115s, std = 41s).

### Cortical local field potential (LFP) recording in conjunction with VSFP imaging in lightly anesthetized mice

LFP activity was recorded from barrel cortex in lightly anesthetized mice (n = 2 males). Animals underwent surgery for the cranial window and a head-bar implant at least a week prior to the recording. At the day of LFP recording, the animals were first put under 2% isoflurane anesthesia for drilling a craniotomy. The glass cover and dental cement above barrel cortex were carefully removed using a dental drill. After the skull was successfully exposed, a small craniotomy with the size of 1 x 2 mm was drilled by the edge of the imaging window. The exposed dura was then covered with a thin layer of Dura Gel (Cambridge Neurotech). For data collection, animals were transferred and head-fixed under the imaging setup. The isoflurane level was lowered to 1%, and the temperature of a heating pad below the mouse was maintained at 37°. A 32-Channel silicon probe with sharpened tip (Part H7b from Cambridge Neurotech) was slowly lowered into the craniotomy at a 45-degree angle to a depth of around 900µm. Acute LFP data were recorded via an RHD USB interface board (Intan Technologies) at 20 kHz. For LFP analyses, the signals across all 32 channels were low-pass filtered with a cut-off frequency of 50 Hz, while the voltage and hemodynamic signals were bandpass filtered between 0.5-50Hz.

### Whisker air puff stimulation in conjunction with VSFP imaging in awake mice

Animals were head-fixed and presented with trains of air puff stimulation to either the right or left side of their face (n = 2 males). Animals underwent the same surgical and habituation procedures as outlined above. Imaging was performed with a 200Hz frame rate, and a single session consisted of a series of 100 trials (50 on each side) of whisker air-puff trains of various frequencies (duty cycle = 50%) that were delivered to the side of the animal’s face. Each trial consisted of 5 s of baseline activity, 6 s of 1 Hz stimulation, 6 s of 2 Hz stimulation, and 3 s of 5 Hz stimulation.

### VSFP image processing

Raw images from both the donor and acceptor cameras were first aligned and registered to the first frame of the green channel for any given trial to perform within trial image registration. The red and green fluorescence signals from the two cameras were analyzed to extract the voltage and hemodynamic signals using the gain-equalization method as previously described (W. Akemann et al., 2012; Carandini et al., 2015; Shimaoka et al., 2018). The gains at the heartbeat frequency were equalized between the two cameras. The gain equalization factors were obtained once per recording for each pixel, using the first trial of the session. The ratio of the two cameras captures FRET signals linked to membrane voltage variations (Carandini et al., 2015) while the sum of the two signals captures large co-fluctuations linked to the hemodynamic response (Shimaoka et al., 2018). To exclude possible residual contamination of the hemodynamics in the ratiometric voltage signal, the sum (hemodynamic) signal was filtered below 5Hz and then scaled by the regression coefficient with the ratiometric signal, then regressed out from the ratiometric voltage signal (hemodynamic signal regression). Hemodynamic signals used for analysis were lowpass filtered below 5Hz to remove physiological noise from the signal (Carandini et al., 2015), except for the hemodynamic signal presented in Figures 2 and 4, which was not filtered for analysis.

**Figure 2.**
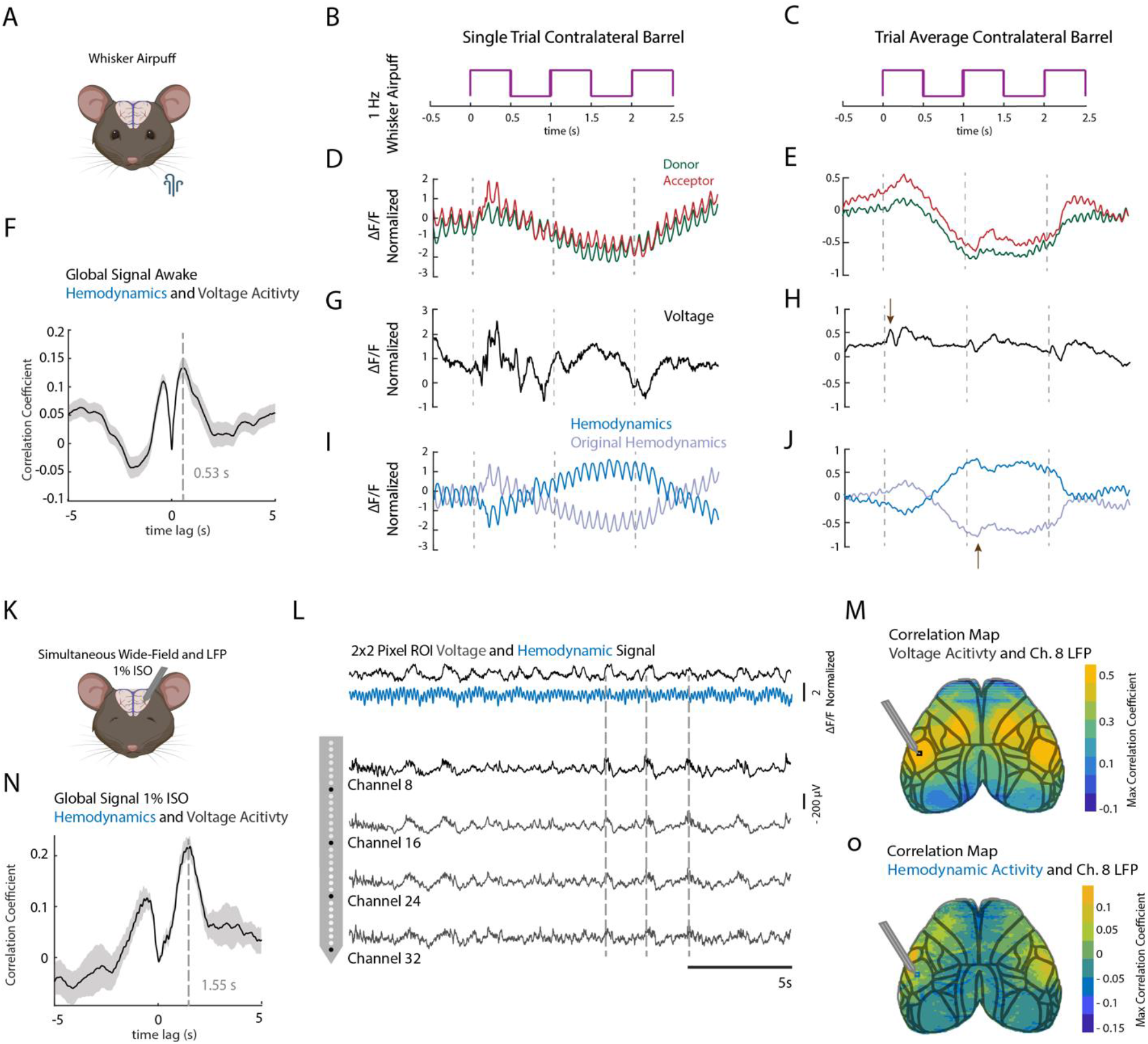
Validation of voltage and hemodynamic signals. **A.** Schematic of mouse representing air puff stimulus delivered to the left side of the face (n=2 mice). **B. C.** Show the time course of the 1Hz air-puff train that was used, duty cycle 50%. **D.** Single trial trace of normalized changes in fluorescence for the raw donor and acceptor channel taken from an ROI over right barrel cortex. Dotted lines represent the onset of the air puff. **E.** Same as in F but for a trial average response (n=100 trials, from 2 mice). **F.** Cross correlation between global hemodynamic and voltage signal. Max correlation of ∼ 0.13 at a lag of 0.53s. Shaded lines represent standard error. **G.** and **H.** The single trial and trial averaged voltage trace obtained from the ratiometric method. Arrow represents the onset of the voltage response. **I.** and **J.** The single trial and trial average hemodynamic response obtained from the ratiometric version. The arrow represents the peak in changes in HbT in response to the air puff stimulation. **K.** Schematic showing the set-up for simultaneous LFP and wide-field imaging. **L.** The top trace (black) shows the average voltage signal from a 2-by-2-pixel ROI next to the LFP electrode as shown to the right on the cortical map (black square). The second row represents the hemodynamic trace for the same ROI. The 4 signal traces below correspond to the positioning of 4 channels on the laminar probe, which was inserted at an angle of 45 degrees into the left barrel cortex at various depths. **M.** The average correlation map shows the correlation between voltage activity and activity from channel 8 of the LFP signal (n = 9 trials at depth ∼1500um). More widely, barrel cortex and an area more anterior in the motor cortex are highly correlated with the LFP trace. **N.** Same as in panel F but for the anesthetized condition. Max correlation is 0.2 at a lag of 1.55s. **O.** Same as in M but for the correlation between hemodynamic activity and LFP. Similar spatial representation as in G, but with a much lower correlation. For all panels that contain the hemodynamic signal, it has been inverted, except for panel I and J.

An increase in the raw hemodynamic signal reflects increases in total hemoglobin which shows up as a decrease in ΔR/R (Shimaoka et al., 2018). To relate these changes to increases in neural activity all hemodynamic signals used for analysis have been sign inverted, to show increases in neural activity and increases in hemodynamic activity both through a positive change in ΔR/R. Photobleaching of Butterfly 1.2 in wide-field imaging conditions is negligible and therefore was not accounted for (W. Akemann et al., 2012).

To perform analyses across trials and mice, the base image used to register within trial frames was aligned to a template cortical map derived from the Allen Brain Mouse Atlas v3 (brain-map.org). A 2D top projection of the annotated Allen Brain Atlas volume was created in MATLAB (MathWorks) and a template image from each mouse was registered to the atlas using an affine transformation computed from two control points that were selected manually: the mid-point at the top of the front portion of dorsal cortex and the base of the cortex at the midline of the retrosplenial cortex. Once all frames were aligned, a mask of the 2D Allen Atlas was created in MATLAB and applied which set all pixels outside of cortex to zero.

### Pupil diameter

To determine the pupil diameter, video frames showcasing the side of the mouse face were first cropped to include only the eye. Pupil diameter was often acquired through the whisker field with whisker motion in front of the pupil limiting the use of existing methods for determining pupil diameter (Reimer et al., 2014; Stringer et al., 2019). A fast pupil detection method using custom-made MATLAB scripts was developed to determine diameter of the pupil frame-by-frame. First, the pixel intensity threshold was set through an interactive user interface to binarize the darkest regions representing the pupil; then the edges of the pupil (which may be composed of multiple objects if whiskers obstruct continuous object isolation) were determined and the distances between all of the edge points was used to calculate the maximum projection across the object(s) representing the pupil diameter. This method is both fast and robust to objects crossing the pupil and creating discontinuities in the ellipse shape of the pupil. Lastly, the pupil diameter trace was smoothed with a 10^th^ order median filter. This removed artifacts such as a portion of the whiskers occluding the pupil across 1-2 frames during diameter detection.

### Orofacial movements

Periods of movement and quiescence were estimated from video capturing the side of the mouse face including: snout, whiskers, and jaw. The onset and offset of movement periods were detected based on average changes in pixel intensity between frames. Orofacial movement signals of each pixel were differentiated and squared. Periods when the frame movement was 2SD above the motion estimate were classified as periods of orofacial movement. Across a single trial, movement periods were classified as points in time where the orofacial movement signal remained above threshold for at least 2 seconds. Rest periods where those periods in which the movement signal remained below threshold for at least 2 seconds.

### Frequency analysis

The time varying power of different frequencies was calculated for all signals using wavelet spectrograms with the Morlet Wavelet. To assess coherence between different pairs of signals we used the built in MATLAB function wcoherence() again using the Morlet Wavelet. The obtained scalograms are susceptible to edge-effect artifacts. These arise from areas where the stretched wavelets extend beyond the edges of the observation interval. Only information within the unshaded region delineated by the white line presents an accurate time-frequency representation of the data. Therefore, all trial-averaged data was obtained by disregarding the data outside of these bounds. For cross-trial comparisons, the magnitude squared coherence values for each single trial was calculated and averaged across trials as a function of frequency. Plots show the mean magnitude squared coherence value, within an ROI or across the whole brain, with error bars representing the standard error.

### Cross-correlation

Unless otherwise indicated all correlation values represent the maximum correlation value obtained from calculating the cross-correlation between two discrete-time sequences using the built in MATLAB function [r,lag] = xcorr(x,y). The cross-correlation measures the similarity between vector x, which is always either the voltage or hemodynamic signal, and shifted (lagged) versions of vector y which in most cases represents the pupil diameter. For figures showing the cross-correlation between voltage and hemodynamic activity we chose x = hemodynamic activity and y = voltage, so that final outputs represent how voltage is shifted in time with respect to hemodynamic activity.

### Global Signal

For whole brain analysis, the global signal was used for both voltage and hemodynamic signals. Global signal represents a timeseries of signal intensity averaged across all pixels in cortex. For each animal and imaging session, global voltage and hemodynamic traces were obtained by averaging the z-scored signal from each pixel located within the cortex while masking pixels outside. Z-scored signals were used to ensure each pixel contributed equally to the calculation of the global signal and that local transients of activity (either noise or physiological) did not bias the global signal calculation.

### ROIs

For ROI specific analysis the signals represent the local averaged signal that represents the shared dynamics of the defined region. ROIs were manually selected once for each animal based on specific areas that show clear signals in the obtained average pupil - voltage activity cross-correlation coefficient map.

### Significance testing

Statistical significance between correlation maps was calculated using Mann-Whitney U-tests which were further Bonferroni corrected to calculate adjusted p-values. Since statistical comparisons were calculated for the extent of the imaging area (10,000 pixels), to guarantee a global error probability at a given threshold (0.05, 0.01, etc.) the significance level for a single test (pixel) was obtained by dividing the global error probability by the number of independent tests. For example, if the p-value was initially significant at p<0.05, the p-value corrected for multiple comparisons would be p<0.05/10000 or p<0.000005.

### Shuffled controls

Correlation maps were tested against chance correlations by randomly selecting the pupil data series along with the voltage and hemodynamic signals for a total of 100 times. Correlation coefficients were calculated on the shuffled data. The original data was then tested against the shuffled data using Mann-Whitney U-tests.

The coherence was also tested against chance by similarly randomly selecting a pupil trace, an orofacial trace, a global voltage trace and a global hemodynamic trace. The magnitude squared coherence for 100 of these new signal pairs was calculated and compared against the original data.

### Code and data availability

All code used for analysis will be made available on our GitHub upon publication. The imaging data presented here will be published on DANDI (https://dandiarchive.org/) in Neural Data Without Borders (NWB) format (https://www.nwb.org/).

### Limitations of the study

Our analysis treats the effects of changes in the vasculature as a time-varying signal that equally influences the donor and acceptor channels. This is a simplification because we know that the absorption spectrum of oxyhemoglobin (HbO) and deoxyhemoglobin (HbR) is different at the donor fluorescence wavelength (542nm) compared to the acceptor fluorescence wavelength (594nm)(Carandini et al., 2015; Ma, Shaik, et al., 2016). While the donor fluorescence wavelength is only slightly less absorbed by HbR the acceptor channel is ∼3 times more strongly absorbed by HbR than by HbT. Given that the ratio of HbO and HbR is constantly changing during ongoing activity, the fractional absorbance of HbO and HbT would differentially affect the donor and acceptor signals. Our analysis method does provide two signals that are separated and follow temporal trajectories that one would expect to see in the characteristic voltage and hemodynamic signals, particularly as can be seen with the whisker air-puff response. We are also not the first to use this sensor to study both voltage and hemodynamics (Carandini et al., 2015; Pisauro, Benucci, & Carandini, 2016; Shimaoka et al., 2018). Yet the degree of separation of the two signals as slower frequencies which are primarily used here for our resting state data remains unknown. More precise measurements of hemodynamic activity could be obtained by using a third camera to record changes at an isosbestic point where the absorption of HbO and HbR is equal (Ma, Shaik, et al., 2016).

Another limitation of our study is the low frame rate for our behavioral tracking (25Hz) which limits the types of orofacial movements we can record, along with the relatively short periods of orofacial movement that we can identify and analyze within our trials. With a mean trial duration of 115s, movement bouts were very short. We therefore were not able to look at the influence of arousal on different periods of motion, despite knowing that the strength of neurovascular coupling changes as a function of the duration of spontaneous movements (Winder et al., 2017). The short nature of our movement periods and the fact that we’re looking at slow frequencies additionally means that what the animal is doing ±4 seconds could contribute to what we were seeing during the movement and rest periods. Future work with longer imaging trials and higher frame rate cameras will allow us to further study the relationship between resting state voltage and hemodynamic activity.

## Results

### Wide-field cortical imaging of voltage activity across dorsal cortex

We used wide-field voltage cortical voltage imaging to record from excitatory neural populations primarily in layer 2/3 over the dorsal surface of the brain during spontaneous behavior in head-fixed mice (see Methods). The VSFP Butterfly protein in transgenic animals was expressed throughout the cortex (Fig. 1A) as previously observed (Carandini et al., 2015; I. Ferezou et al., 2007; Madisen et al., 2015; Ratzlaff & Grinvald, 1991). Wide-field optical signals are very sensitive to physiological variables such as blood flow, blood oxygenation, and intrinsic autofluorescence (Ma, Shaik, et al., 2016; Pisauro et al., 2016). A ratiometric equalization technique (Carandini et al., 2015) was used to compare the two combined fluorescent signals from both the acceptor and donor channel of VSFP Butterfly to estimate average local membrane voltages separately from the hemodynamic signals (Fig. 1C - D). Video of the face was recorded to track changes in orofacial movements and pupil diameter (Fig. 1E). Our algorithm picks up changes in orofacial movements when there is a noticeable differences between individual frames induced by movements of jaw, tongue or nose (Fig. 1F). The pink boxes highlight which regions in the frame change which show up as a change in orofacial movement using our method.

To validate the calculated voltage and hemodynamic signals we tracked the response to a whisker air puff train delivered to the left side of the face (Fig. 2A, n= 2 awake mice). The air puff stimulus consisted of a 6 s long 1 Hz train, with a duty cycle of 50% (Fig. 2B – C). Single trial activity traces represent the average activity from an ROI in right barrel cortex while the trial average is the average activity across both mice in right barrel cortex (n = 100 trials). Following the onset of the first air-puff stimulation both green and red fluorescence channels experience a slow decrease superimposed on large pulsations associated with heart beats (Fig. 2D), when averaging across multiple stimulus presentations the large pulsations which are not phase locked to the stimulus presentation are averaged out (Fig. 2E) leaving a clear stimulus-triggered hemodynamic response (common signal in green and red fluorescence). These slow responses to the stimuli do not reflect neural activity as responses to whisker air puff stimulation in awake mice have an onset latency of only a few ms (Petersen, 2019). Using the ratiometric method we were able to resolve a similarly fast clear bi-phasic voltage response that is triggered by the onset of the air puff stimulation (Fig. 2H) The hemodynamic signal obtained from the ratiometric method contained the characteristics that would be expected from changes in blood volume and oxygenation that occur in response to the onset of the stimulus. For a single trial a strong oscillatory component overlaid on top of a slow hemodynamic response was present (Fig 2I). When averaging across trials a slow response peaking at ∼1.2 seconds after stimulus onset remained (Fig. 2J). This is consistent with an increase in total hemoglobin that resulted in a decrease in reflectance caused by arterial and capillary dilations (Winder et al., 2017). Note that we are showing a hemodynamic signal that has been sign-inverted to show both increases in voltage activity and increases in hemodynamic activity with positive signals (dark blue traces, Fig. 2I and J). When comparing these two global signals directly we observed a bimodal response. In the anesthetized relative to the awake condition, we observe an increase in the maximum correlation (0.2, anesthetized versus 0.1, awake) and an increased latency for the anesthetized state relative to the awake condition (1.55s versus 0.53s), as would be expected (Fig. 2F and 2N).

Furthermore, we looked to determine the relationship between the voltage and hemodynamic imaging signals as they relate to electrophysiological voltage activity across cortical layers. To this end we recorded laminar LFP signals simultaneously with voltage imaging (Fig. 2k). A linear array of recording sites of a 32-channel silicon electrode was placed to span multiple layers in the barrel cortex in lightly anesthetized mice (Fig. 1L). LFP waveforms showed a clear similarity with voltage imaging traces determined with the ratiometric method from our raw signal from adjacent areas of cortex, while the hemodynamic signal contained the same underlying slow frequency oscillations but was dominated by larger heartbeat pulsations (Fig. 2L). We found barrel cortex LFP was also strongly coupled to voltage imaging traces in contralateral barrel cortex as well as ipsilateral and contralateral motor cortex with a maximum correlation of 0.5 (Fig. 2M), revealing a sensorimotor network of co-activity under light isoflurane anesthesia. Hemodynamic activity on the other hand was much less correlated, with a maximum of 0.1 (Fig. 2O). These findings indicate a clear relationship between LFP activity and layer 2/3 voltage imaging, though voltage imaging is showing signal averages across 0.1 x 0.1 mm areas of cortex, and contributions from deeper layers likely differ between the two methods.

### Pupil diameter is tightly coupled to global cortical signal changes in voltage and to a lesser extent to hemodynamics

First, we were interested to determine whether cortical signal changes might globally fluctuate with arousal as measured by pupil diameter. When plotting the global voltage signal and pupil diameter activity together, a strong similarity in the resulting waveform is readily apparent (Fig. 3A), with a possible delay between voltage and pupil signal. To quantify and examine the variability of time lag between the pupil diameter and the global voltage, a cross-correlation between the global signal and the pupil diameter was calculated (see Methods). A lag was not observed when data from all mice were averaged. What remained consistent, however, was the strong correlation between the two signals with a peak correlation of ∼0.5 (Fig 3C). These results indicate a strong relation between average cortical activation and arousal as measured by pupil diameter. Previous work imaging noradrenergic and cholinergic axons in layer 1 of visual cortex showed a delay of 0.5-1 s between activation of these arousal-related neuromodulatory systems and pupil diameter in rats (Reimer et al., 2016), suggesting that global cortical depolarization increases we observed have a similar lag to neuromodulation, given that our lag with relation to pupil diameter was close to zero.

**Figure 3.**
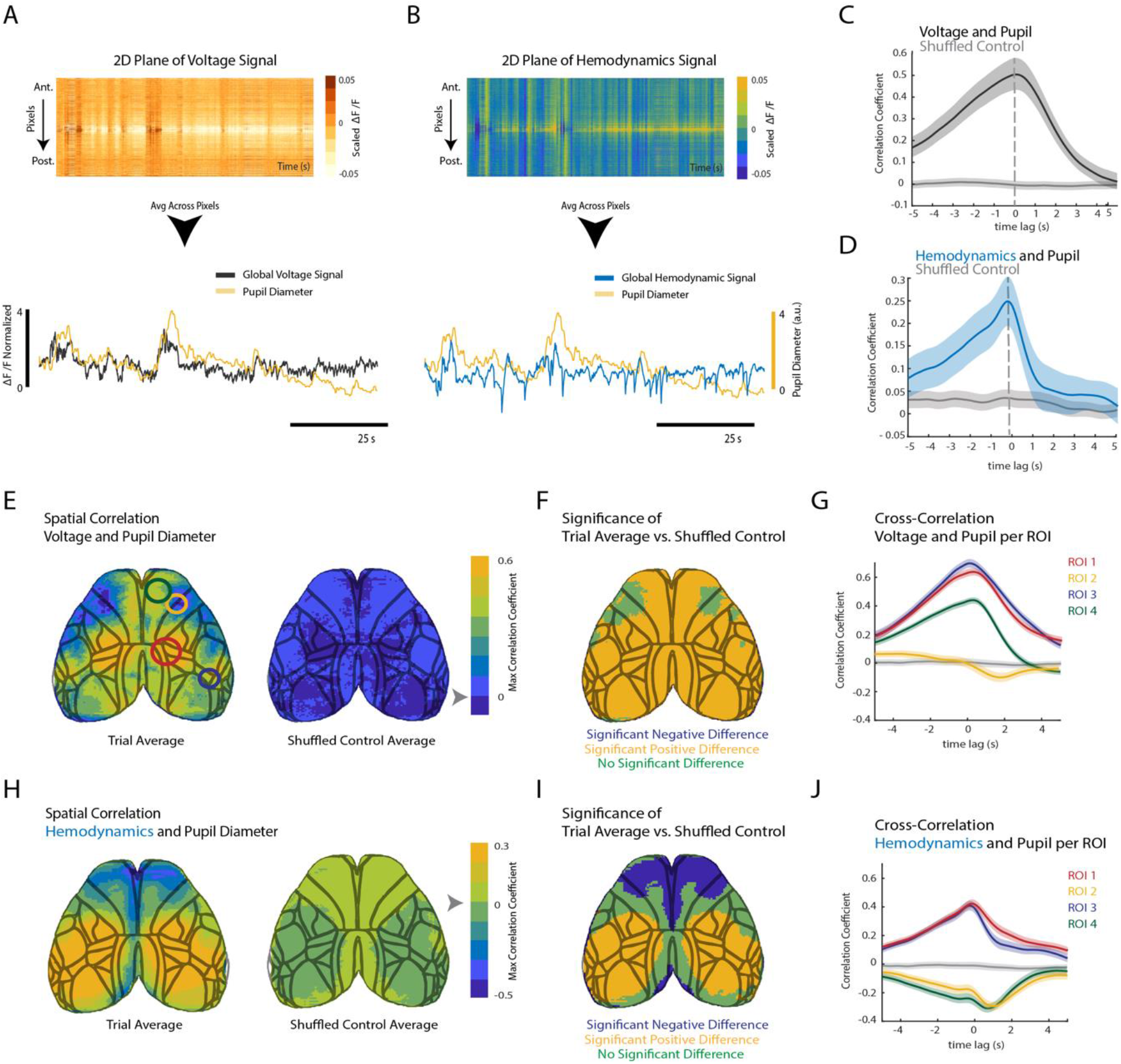
Region dependent coupling of pupil diameter to voltage and hemodynamic activity across functional cortical areas. **A.** All pixels from the cortical surface are laid out in a 2D plane such that the changes in voltage activity for each pixel (y-axis) is represented as a function of time (x-axis). The global signal is obtained by taking the z-score of each pixel and averaging across all pixels. The global voltage signal (yellow) fluctuates in a similar manner to pupil diameter (grey). **B.** Same as in panel A but for cortical hemodynamics signal. A relationship between changes in pupil diameter (grey) and subsequent changes in global hemodynamics (blue) can be observed. Correlation between pupil diameter and global signal is consistent across trials and animals. Where shaded area represents the standard error. **C.** Average cross-correlation between voltage and pupil diameter results in a high correlation of 0.5 centered at ∼0s. **D.** Average cross-correlation between hemodynamics and pupil diameter also shows a positive coupling. With max correlation of 0.25 also centered at ∼0s. Panels C-D show average across n=65 trials for 5 mice. Grey trace indicates shuffled control. *The shaded area in all figures represents standard error.* **E.** The panel on left represents the average correlation map obtained from all 5 animals (n = 65 trials) between voltage traces and pupil diameter. Strong correlations between voltage activity and pupil diameter can be observed within the medial sensory-motor cortical areas as well as more laterally in the secondary sensory/ auditory cortices (ROI 1 and 3). There exists also a slight negative correlation with pupil diameter in the more lateral areas of motor cortex (ROI 2). The panel on the right represents the correlation map obtained between 100 randomly selected imaging trials and 100 randomly selected pupil video files. There are no areas that are strongly correlated by chance. **F.** This map shows the significantly different areas between the trial average map and the shuffled control at p < 0.05 adjusted for multiple comparisons with Bonferroni Correction (BF). Blue and Yellow represent areas that have a significant negative and positive difference to the control respectively, while green is areas with no significant difference. The map highlights significance across most areas, including ROIs 1,3 and 4. ROI 2 which corresponds to the more lateral area of motor cortex was not statistically different when comparing to the average map. **G.** Average cross-correlation between voltage and pupil diameter per ROI shows region dependent coupling. **H.** Panel on the left shows the average coupling between hemodynamic signal and pupil from cross-correlations shows similar positive correlations as seen in voltage activity (ROI 1 and 3). Most notably however is the reduction in correlation, now with a peak slightly above 0.3 as well as the stronger negative correlation of -0.4 that spans primary and secondary motor areas (ROI 2 and 4). Right shows the same as panel C but for hemodynamics. **I.** Same as in panel F but for the relationship between the average trial hemodynamics map and the shuffled control. **J.** Same as in panel G but for hemodynamics and pupil diameter. Note the difference in response for ROI 4 and the different lags for negative peaks. Grey line indicates the shuffled control for ROI 1. Controls for all other ROIs were nearly identical and therefore are not plotted. For all panels that contain the hemodynamic signal, it has been inverted and lowpass filtered at 5 Hz. For a similar analysis done using the orofacial movement trace see extended data Figure 3-1.

**Figure 3-1.**
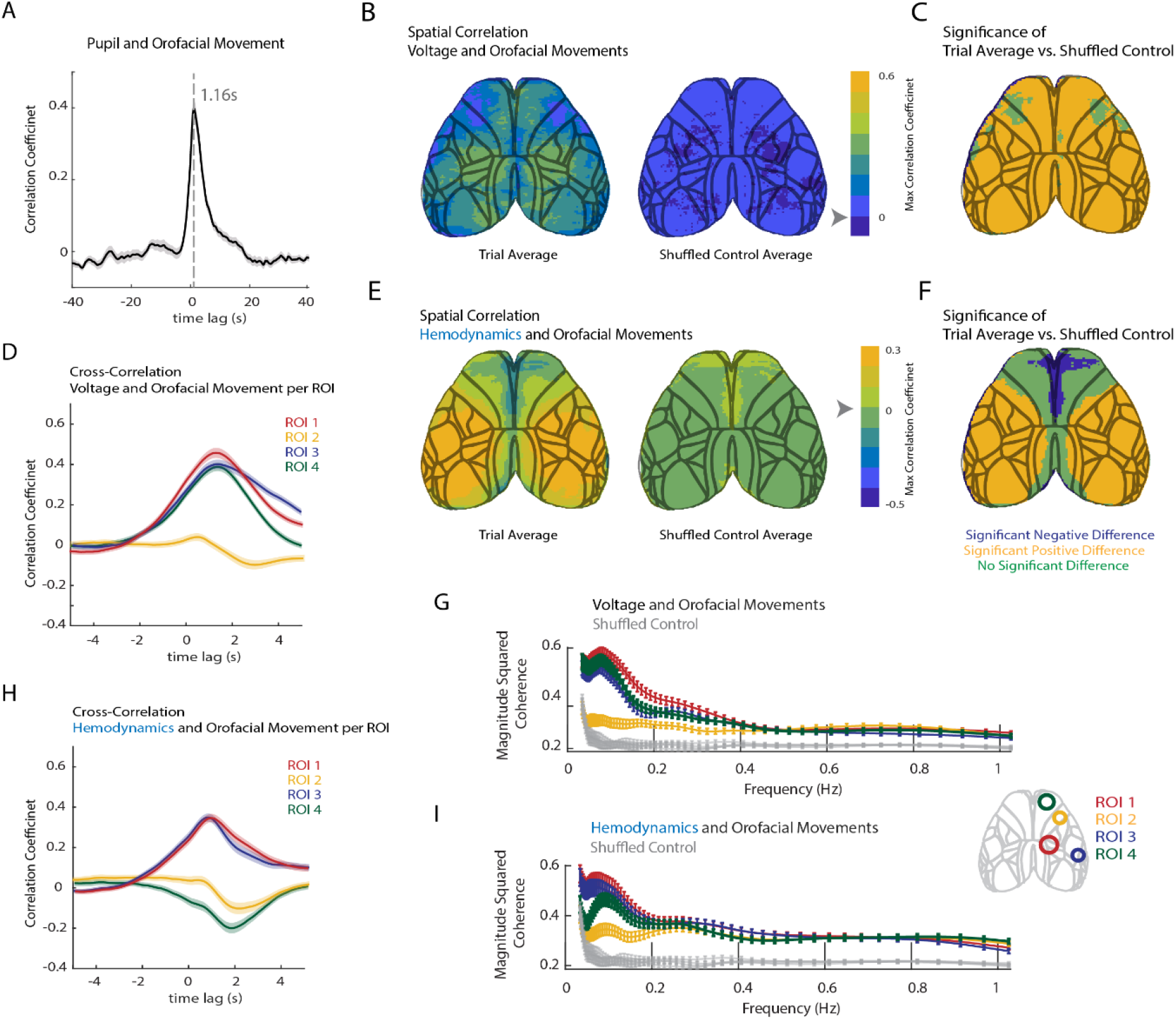
Region dependent coupling of orofacial movements to voltage and hemodynamic activity across functional cortical areas. **A.** Trial average cross correlation between pupil diameter changes and orofacial movements. Max correlation of ∼0.4 at a lag of 1.16s. **B.,C.,E.,F.** Same type of spatial cross correlation maps as in Figure 3 panels E,F,H,I but correlating now changes in activity to changes in orofacial movements. **D., H.,** Same cross correlations per ROI as shown in Figure 3 panel G and J but now with relationship to changes in orofacial movements. **G.** Magnitude squared coherence between voltage activity and orofacial movements across 4 different ROIs. **I.** Same as in G but for hemodynamic activity and orofacial movements. Error bars represent standard error. For all panels that contain the hemodynamic signal, it has been inverted and lowpass filtered at 5 Hz.

Global changes in hemodynamic activity were far less correlated to changes in pupil diameter than global voltage was (Fig. 3B). The peak correlation between hemodynamics and pupil diameter was determined by fixing each trace of global hemodynamics and shifting the pupil trace. When shifted traces yielding maximal peak correlations were averaged, a similar temporal relationship between hemodynamics and pupil diameter was observed with a maximum correlation of ∼0.25 (Fig. 3D), suggesting that global hemodynamic changes were also correlated to changes in pupil diameter but to a lesser degree. This result matches what was previously found for the “global” hemodynamic signal in primary visual cortex in awake mice using intrinsic optical imaging where an average net zero time lag with a maximum correlation of 0.36 was observed (Pisauro et al., 2016). In contrast more recent work looking at hemodynamic activity and pupil diameter in somatosensory cortex for awake mice found a peak correlation of -0.37 ± 0.25 with a delay of 1.3 seconds (Turner, Gheres, Proctor, & Drew, 2020) which implies that there is an inverse relationship between these two signals during resting state. These authors further showed that the orientation and degree of correlation was state dependent. Overall results suggest that the correlation between the two signals is spatially variant and perhaps dependent on behavioral state.

### Cortical activity and pupil diameter are differentially coupled across functional cortical areas

Our data analysis so far supports a link between changes in arousal as measured by pupil diameter and changes in global cortical voltage and to a lesser extent the hemodynamic signal. To probe for a differential contribution of different cortical areas on the observed correlations, we calculated correlation maps between pupil diameter and changes in cortical voltage and hemodynamics signal at each pixel. Spanning dorsal cortex, we identified 4 specific ROIs (Fig 1B) that displayed a distinct relationship between voltage and arousal bilaterally. We found that the maximum correlation coefficients between voltage and pupil diameter were highest around medial somatosensory and secondary somatosensory and auditory cortex (Fig. 3E, max correlation 0.6). Voltage activity of medial aspects of frontal cortex corresponding to parts of secondary motor and anterior cingulate cortex was also strongly coupled with pupil diameter (Fig. 3E ROIs 1,3 and 4). In contrast, there was on average zero correlation between pupil diameter and voltage signal in more lateral areas of motor cortex (Fig. 3E, ROI 2). Overall, however, positive correlations predominated, explaining the positive correlation of global voltage signal with pupil diameter. To assess the statistical significance of observed correlations with voltage, we compared average correlation maps across all trials (n = 65) to a shuffled control (n =100), where voltage traces and pupil diameter traces were taken from different trials. This analysis revealed a significant correlation between voltage activity and pupil diameter for ROIs 1,3 and 4 (Fig. 3F). A net zero correlation was observed in ROI 2.

We observed a much weaker relationship between pupil diameter and global hemodynamics. However, it is possible that the correlation between pupil diameter and hemodynamics could be more spatially heterogenous. Like voltage, hemodynamics also showed a significant positive correlation with pupil diameter in barrel cortex and more medial somatosensory areas (Fig. 3H). The maximal correlation coefficient was lower with 0.2 as opposed to 0.6 for voltage, however. Hemodynamic activity in contrast showed larger and stronger negative maximal correlation peaks than voltage with pupil diameter in frontal medial cortex (Fig. 3H, blue areas). We assessed the statistical significance of observed correlations with hemodynamics, using a shuffled control (n =100), where hemodynamic traces and pupil diameter traces were taken from different trials. This analysis revealed a significant correlation between hemodynamic activity and pupil diameter for all 4 ROIs, with ROI 1 and 3 being positively different and ROI 2 and 4 negatively different (Fig. 3I). We plotted the cross-correlation function for each ROI to determine the detailed time relationship between pupil diameter changes compared to voltage or hemodynamic activity changes (Fig. 3G,J). We found that all positive correlations had their peak with a delay near zero, indicating no average time shift between voltage, hemodynamics, and pupil diameter changes. In contrast, negative correlations between hemodynamics and pupil diameter had a peak correlation at a delay of about 1s, indicating that increases in pupil diameter preceded decreases in hemodynamic activity in these ROIs.

Another widely used metric for tracking changes in arousal state, and for breaking up awake resting imaging data into different behavioral states, is to use the self-generated movements as a proxy for arousal (Drew, Winder, & Zhang, 2019; Winder et al., 2017). We wanted to see how the correlations would change if we used spontaneous orofacial movements rather than pupil diameter as our metric of arousal. As expected, these two signals were correlated with each other, with changes in orofacial movements preceding changes in pupil diameter by 1.16 seconds (Fig. 3-1A). Spontaneous behavioral changes are known to drive neural activity through various mechanisms, resulting in temporally varying activations of multiple brain regions (Drew et al., 2019). In agreement with such variable neural activation, we observed a lower spatial correlation with voltage activity than we found with pupil diameter (Fig. 3-1B, max of 0.4). But we still found significant differences related to the shuffled control across all cortical areas (Fig. 3-1C). In contrast the hemodynamic maximal spatial correlation was not noticeably different than what we observed with pupil diameter (Fig. 3-1E). However, we did observe a decrease in the number of pixels in prefrontal cortex that were negatively correlated (Fig. 3-1F).

Overall, voltage correlation with pupil diameter showed widespread significant positive correlations including medial prefrontal areas, whereas hemodynamics showed a negative correlation in the same frontal network while maintaining positive correlations in more posterior areas. We also observed similar patterns when using orofacial movements rather than pupil diameter as a proxy for arousal. It is important to note, however, that changes in pupil diameter also occur in the absence of orofacial movements.

### Low frequency dependence for pupil diameter, cortical activity, and neurovascular coupling

Next, we sought to determine whether the observed correlations depend on coupling at specific signal frequencies. Visual inspection of both voltage and hemodynamic signals from an ROI in relation to changes in pupil diameter suggests that the observed coupling was driven by slow frequency components present in each of the signals (Fig 1D). Looking at the average amplitude power spectrum for the global voltage and hemodynamic signals up to 14Hz we see that most of the power is concentrated at the lower frequencies (Fig. 4 A). With the exception of the hemodynamic signal which has a clear peak centered at 12 Hz which coincides with the heartbeat frequency of awake mice (Fig. 4B) and reflects the hemodynamic change related to heartbeat.

**Figure 4.**
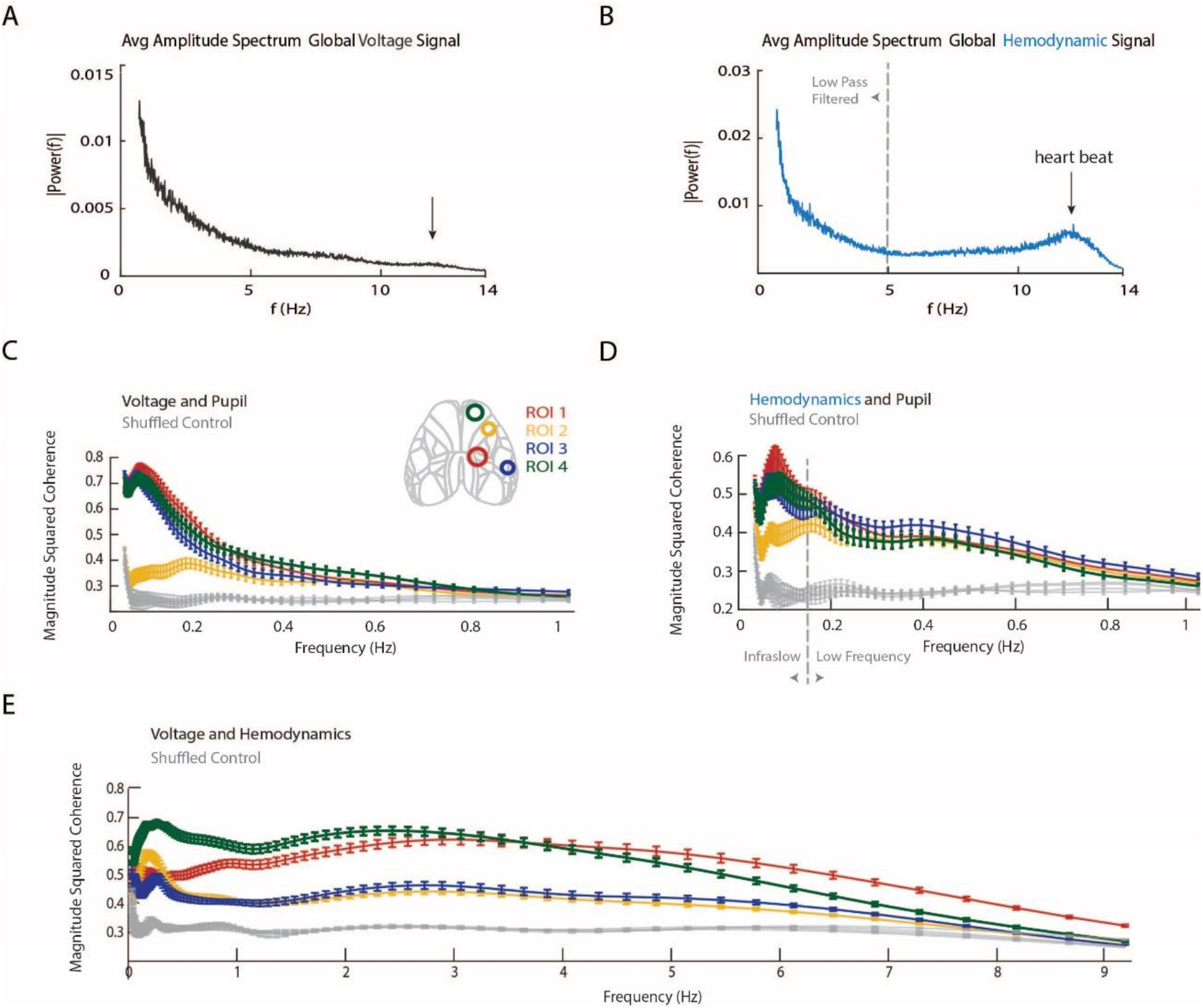
Region dependent frequency coupling of pupil diameter to voltage and hemodynamic activity. **A.** Average amplitude spectrum from the global voltage signal (n = 65 trials). Signal shown is truncated between 0.7 – 14 Hz to improve visibility of the higher frequencies. **B.** Same as in A but for hemodynamic activity. Clear peak can be seen centered around 12 Hz corresponding to heat beat. **C.** Plot shows the average magnitude squared coherence for all trials for pupil and voltage within the cone of influence. **D.** Average magnitude squared coherence for all trials for pupil and hemodynamics within the cone of influence. Allen Atlas is for referencing ROI location. **E.** The magnitude squared coherence between hemodynamics and voltage across all trials within the cone of influence. Each line represents the average values from each ROI with bars indicating the standard error. For panels E,F and G each line represents the average values from each ROI with the bars indicating the standard error (n = 65 trials, 5 mice). Due to edge effects, less credence is given to areas of apparent high coherence that are outside or overlap with the cone of influence. Plots therefore disregard the first and last 20 seconds. The 4 grey lines for panel C-E represent the average magnitude coherence (n = 100) obtained by comparing a randomly selected hemodynamic traces with a randomly selected voltage trace from different trials for each ROI. The error bars represent standard error. For all panels that contain the hemodynamic signal, it has been inverted but no lowpass filer has been applied.

We further analyzed frequency dependence of voltage and hemodynamic activity related to arousal using wavelet coherence analysis. When looking at voltage and pupil, these two signals were most coherent in the low frequency bands below 0.25Hz for ROIs 1,3 and 4 (Fig. 4D), except for ROI 2 that showed a zero-overall coherence between voltage and pupil (Fig. 4C). The three ROIs where voltage positively correlated with pupil diameter showed very similar coherence with a peak magnitude squared coherence of ∼0.8 just below 0.1 Hz. Hemodynamics showed a similar frequency relationship to pupil diameter as voltage, though the peak below 0.1 Hz was smaller with a peak ∼0.6, while the coherence between 0.2 and 0.6 Hz was of similar amplitude of 0.4. Higher frequencies not plotted showed no significant coherence when compared to the shuffled control (Fig 4C and D). Coherograms between voltage and hemodynamics (Fig. 4E), showed a peak at low frequencies centered around 0.2 Hz. In addition, voltage and hemodynamics showed a plateau of strong coherence between ∼1.5 to ∼6 Hz (Fig. 4E). A shuffle control (Fig. 4 E, grey lines) showed no such coherence and indicated that the coherence between voltage and hemodynamics was highly significant up to 9 Hz. This finding indicates that voltage and hemodynamics share higher frequency dynamics that are not related to pupil diameter changes.

### Patterns of cortical coupling to pupil diameter are dependent on behavioral state

Previous studies have described large effects of behavioral state on membrane voltage, hemodynamics, and neurovascular coupling across multiple brain regions (I. Ferezou et al., 2007; Poulet & Petersen, 2008; Shimaoka et al., 2018; Winder et al., 2017). Brain activity varies with arousal, attention, and behavior (Winder et al., 2017). The quiescent periods between bouts of exploratory behaviors in the absence of an overt task, termed “resting periods,” are known to also exhibit stereotyped patterns of brain activity. In primary sensory areas these fluctuations in brain activity are tracked by small fluctuations in pupil diameter (Reimer et al., 2014). To characterize the effects of behavior on the coupling between cortical voltage and hemodynamic activity and pupil diameter, we separated cortical activity between periods of movement and rest. Movement periods were determined from a video of the face as periods of spontaneous orofacial movements. These orofacial movements consisted of changes in whisker, jaw, or snout position (Fig. 1F) and occurred in distinct bouts (Fig. 5A). Pronounced pupil diameter changes were frequently associated with periods of movement, whereas such changes were more modest, but still present, in the absence of movement (Fig. 3-1A, Fig. 5A).

**Figure 5.**
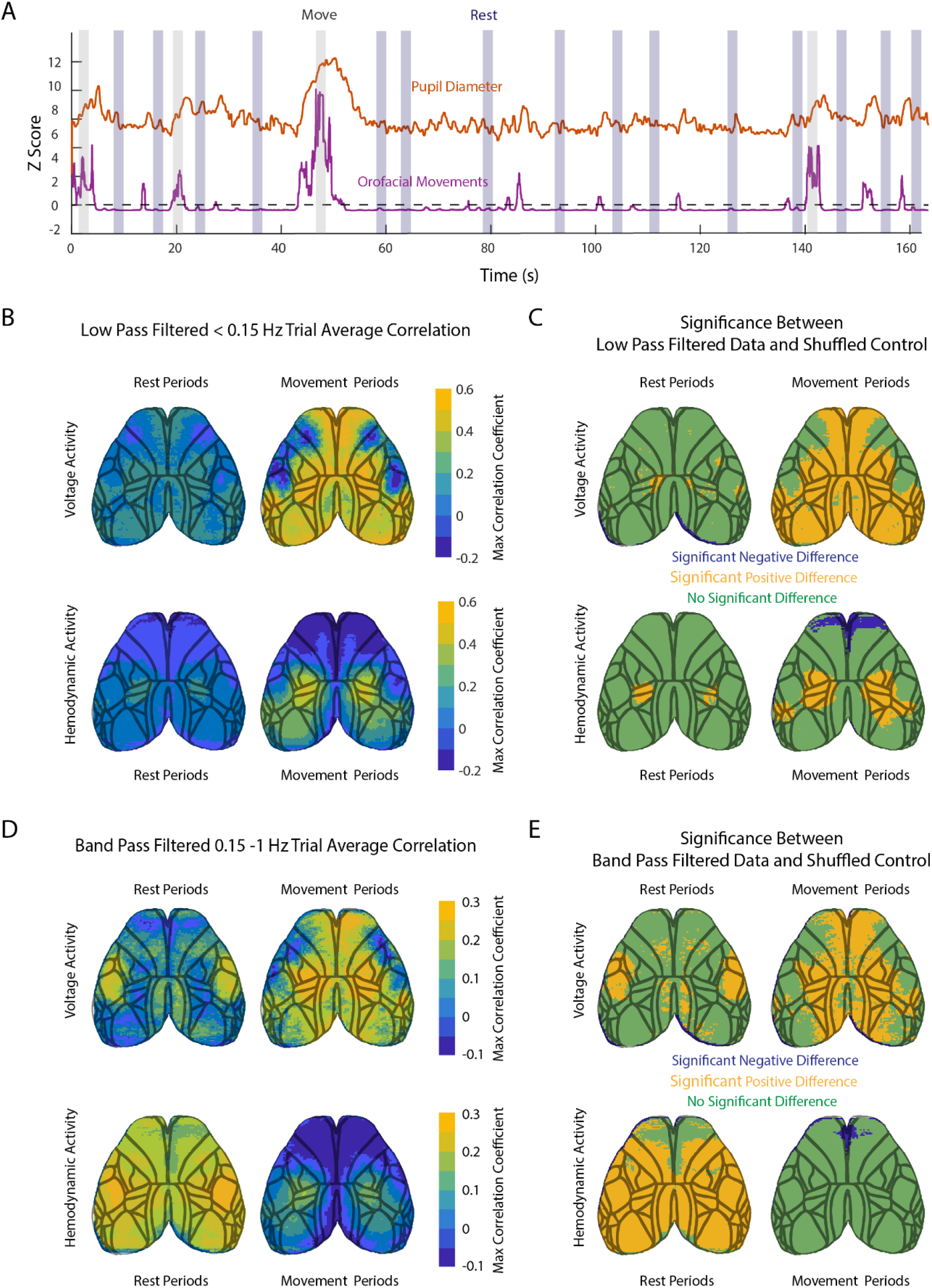
Patterns of cortical coupling to pupil diameter are dependent on behavioral state. **A.** Example trial showing pupil diameter (red) and orofacial movement (purple). Example movement periods (shaded gray) were classified based on the movement signal remaining above the threshold (dashed line) for at least 2s. Periods of rest were instead identified when the movement signal remained below the threshold for at least 2 seconds. **B.** Top left shows the average correlation map (n = 65 trials) for cortical voltage activity (lowpass filtered <0.15Hz) and pupil diameter for periods of rest. Top right shows the voltage correlation map during periods of facial movement. The bottom panels show the same analysis for hemodynamic activity. **C.** Statistical map showing significantly different areas between what’s displayed in panel B with a shuffled control, adjusted for multiple comparisons. **D.** Same activity and behavior trials as in C but now bandpass filtered (0.15-1 Hz). **E.** Same statistical significance as shown in panel C but for band pass filtered data in E. For panels C and E the areas labeled in blue and yellow represent areas that have a significant negative and positive difference to the control respectively, while green is areas with no significant difference. For all panels that contain the hemodynamic signal, it has been inverted and lowpass filtered at 5 Hz.

The data was further split into two frequency bands, a low passed (LP) infraslow signal <0.15Hz and a band passed (BP) 0.15 – 1 Hz (Fig 4D) slow activity signal. These two frequency bands displayed distinct coherence with pupil diameter (Fig. 4C, D) and are also within the range of frequencies that can be resolved with fMRI.

Our analysis revealed substantial differences between cortical voltage and hemodynamic activity coupling to pupil diameter dependent on the presence of spontaneous movements. When animals were resting, correlations between pupil and voltage or hemodynamic activity were near-absent in most cortical areas for activity below 0.15Hz. The strongest correlations that were observed during rest for this low frequency range were at 0.2 (Fig. 5B and 5D, left panels) and localized to primary somatosensory areas close to the midline and posterior parietal cortex.

In the presence of orofacial movement, large areas of cortex showed correlations between voltage and pupil diameter below 0.15 Hz, while the correlations between hemodynamics and pupil diameter were much more restricted to medial somatosensory areas (Fig. 5B, right panels). The highest correlations were confined to the upper and lower limb somatosensory areas. Statistical comparison between rest and movement periods compared to a shuffled control revealed a significant influence of medial sensory areas during both behavioral states for voltage, with movement periods driving also significant correlations in prefrontal and more anterior spanning sensory, visual, and auditory cortices (Fig 5C, top panels). In comparison hemodynamic signal correlations were most significant during movement, with strong positive changes observed in medial sensory cortex and significant negative changes in prefrontal cortex (Fig 5C, bottom panels).

Cortical activity signals at frequencies between 0.15 and 1 Hz showed a substantially different relationship with orofacial movements than the infraslow signal. Here, a pronounced widespread correlation was found between hemodynamics and pupil diameter at rest (Fig. 5D, bottom left), which was largely absent during movement (Fig. 5D, bottom right). A relation between voltage and pupil diameter was also present in this frequency band, but this was spatially limited to barrel cortex (Fig. 5D and 5E, top left). This relationship may be due to whisker movement during rest which we were not able to detect with our code due to the few numbers of pixels that contain whiskers. Changes in these pixels did not induce large enough changes in frame-by-frame variance to be detected as a movement bout. Interestingly, during orofacial movement periods, this relationship disappeared in barrel cortex, but instead a different area more medially now correlated strongly with pupil diameter (Fig. 5D and 5E, top right). These findings indicate the presence of two distinct functional relations between pupil diameter and cortical voltage activity between periods of absence or presence of orofacial movements. In marked contrast, hemodynamic activity was more exclusively and globally related to pupil diameter at rest, and only weekly during movement.

Overall, these data reveal a clear dissociation between how the hemodynamic signal relates to pupil diameter in comparison to brain voltage activity. Further, the relationship for both signals is distinctly different for infraslow frequencies than frequencies between 0.15 and 1 Hz.

## Discussion

In this work we characterized the temporal and spatial components of cortical coupling to arousal as measured through changes in pupil diameter by looking at both changes in voltage activity and hemodynamics across dorsal cortex.

### Cortical networks linking spontaneous fluctuations to changes in arousal

Spontaneous fluctuations in the global brain state have been described to encode changes in cognitive variables such as changes in arousal state (Jacobs, Steinmetz, Peters, Carandini, & Harris, 2020; Pisauro et al., 2016) and are known to contribute to the apparent noisiness of sensory responses at both the neural and behavioral levels (McGinley et al., 2015). While the global signal is often considered a nuisance variable in fMRI imaging, we found that the global signal for voltage activity, and to a lesser extent hemodynamics, was coupled throughout cortex to changes in arousal as measured by pupil diameter (Fig. 3) (Fox et al., 2008; Macey, Macey, Kumar, & Harper, 2003). Furthermore, our results show that arousal influences distinct cortical regions for both voltage and hemodynamic activity. For both signals there is a broad positive correlation across most sensory-motor cortices, extending posterior all the way to primary visual cortex. The strongest positive correlations stem from the medial sensory-motor areas corresponding to the upper and lower limbs, posterior partial cortex, along with more lateral regions in secondary sensory/auditory cortices. In contrast in frontal motor/cognitive cortical areas, we observe differences between the voltage and the hemodynamic signals. For the voltage activity we see a strong positive correlation along the midline in secondary motor cortex which corresponds broadly to eye/eyelid movements as well as the primary motor representations for the rostral and caudal forelimb. In contrast the hemodynamic activity shows a small net negative correlation to pupil diameter across most of secondary motor cortex including the midline. Thus, removal of global signal during pre-processing should be carefully considered because its influence is spatially variant and serves as a covariate for changes in arousal or attention (K. Murphy, Birn, Handwerker, Jones, & Bandettini, 2009; Xu et al., 2018).

The largest network difference between the voltage and hemodynamic signals, when related to changes in arousal, is observed in the pre-frontal cortex. Voltage activity in pre-frontal cortex is positively correlated with arousal while there is a net negative correlation observed in these same areas for hemodynamic activity. In human BOLD-fMRI research pupil-related activations are known to influence cortical areas that regulate selective attention, salience, and decision-making (DiNuzzo et al., 2019). To what degree resting-state networks assessed with fMRI reflect electrical neuronal population activity, and whether electrical activity shows the same relationship to behavioral and cognitive variables as hemodynamics is not well understood. Our results highlight the possibility that arousal has differential influences on pre-frontal cortex during resting state for hemodynamics and voltage activity which can have subsequent implications for how resting state networks are identified and interpreted.

### Temporal profiles of resting state signals

The spectral decomposition of our observed arousal signals fluctuates across both fast and slow timescales. We found that arousal changes as reflected by fluctuations in pupil diameter were differentially coupled to voltage and hemodynamic activity across distinct frequencies (Fig. 4). The spectral decomposition revealed the strong coupling between pupil with voltage and hemodynamic activity up to 0.2 Hz. This is particularly relevant for BOLD-fMRI which typically cannot capture frequencies > 0.5 -1 Hz due to limitations of the imaging system and inherent low-pass filter properties of the vasculature (Kim, Richter, & Uǧurbil, 1997).The slow fluctuations on the order of seconds that were observed in both voltage and hemodynamic activity therefore track the slow fluctuations in arousal (Reimer et al., 2014). Distinct time courses for each frequency band could represent state fluctuations between periods of low and high arousal. Low arousal represented by pupil constriction was characterized by an increased low-frequency oscillations and higher ensemble correlations (Chan, Mohajerani, LeDue, Wang, & Murphy, 2015; Reimer et al., 2014). Pupil dilation, representing an increase in arousal and a subsequent increase in locomotion results in a suppression of a low frequency (<4Hz) band (McGinley et al., 2015).

### Patterns of cortical coupling to pupil diameter are dependent on behavioral state

Expanding on initial attempts to link arousal related to voltage activity across cortex for periods of locomotion, we demonstrate that pupil dynamics were coupled to neural activity also in the absence of locomotion (Fig 6) (Shimaoka et al., 2018; Vinck et al., 2015). While there is no locomotion in our experiments, these head-fixed mice do undergo periods of complete stillness/rest and periods of prolonged uninstructed orofacial movements. Such orofacial movements have been shown with wide-field calcium imaging to be associated with strong neural activity changes over complex cortical networks (Musall, Kaufman, Juavinett, Gluf, & Churchland, 2019). We find that like locomotion, orofacial movement is associated with periods of heightened arousal indicated by increased pupil diameter (Fig. 3-1A, 5A). Changes in arousal as measured by locomotion are known to lead to increased neural activity in motor areas, retrosplenial cortex, and sensory cortices (Clancy, Orsolic, & Mrsic-Flogel, 2019). We found that in a similar manner periods of uninstructed orofacial movements resulted in increases in correlation between voltage and pupil in medial sensory-motor areas and decreased correlations in lateral motor and sensory areas.

**Figure 6.**
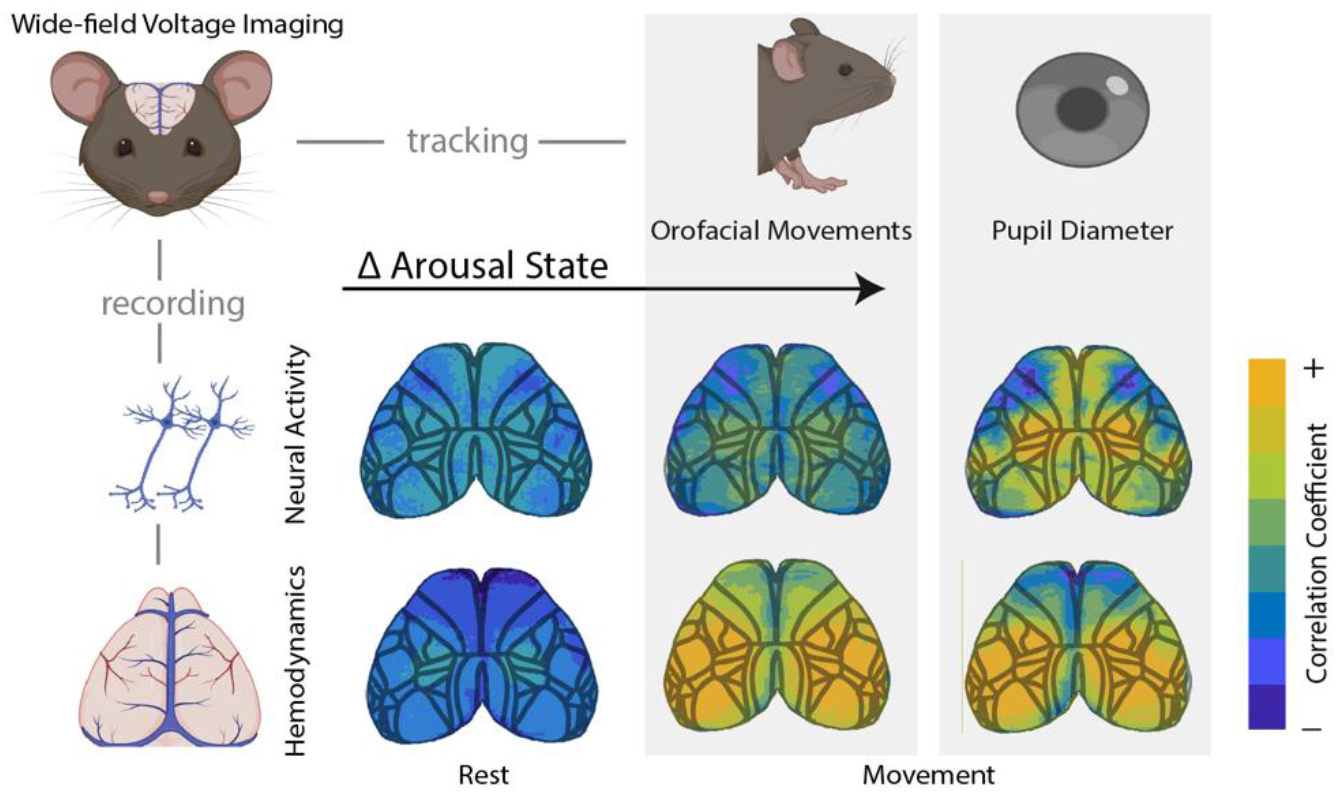
Summary Figure. With wide-field voltage imaging we were able to track changes in both neural activity and hemodynamics across dorsal cortex in waking mice. Changes in arousal state as measured by pupil diameter are differentially correlated with different cortical areas during rest versus during periods of orofacial movement. As a measure of behavioral state, orofacial movements can be used to separate “resting state” data into true rest and movement periods. This reveals that cortical coupling to pupil diameter is dependent on the behavioral state of the animal.

Additionally, we find that activity from movement periods drives the correlations we see during the trial average response, when not accounting for differences in the behavioral state of the animal (Fig. 3). Movement periods result in the strongest correlations seen in the upper and lower limb somatosensory areas. This activation pattern corresponds to increases in limb movements that accompany periods of fidgeting (Winder et al., 2017). Looking at correlations in distinct frequency bands results in different spatial representations. Note that the differences between resting state activity for low pass and band-pass filtered activity is likely explained by the fact that whisker deflections in the absence of other overt movements are not detected by our method.

The largest difference we observe across states when comparing the voltage and hemodynamic correlation maps with pupil diameter was in prefrontal cortex. Previous work has revealed some intricacies between neurovascular coupling and the behavioral state of the animal as a function of explicit sensory cortical areas (Cardoso, Lima, Sirotin, & Das, 2019; Huo, Smith, & Drew, 2014; Winder et al., 2017). One study showed that in prefrontal cortex during voluntary locomotion there was a decrease in cerebral blood volume (CBV) despite recording increases in multiunit activity. Our voltage imaging results complement this by finding a similar dissociation between our hemodynamic signal and our voltage activity map in prefrontal cortex during periods of increased orofacial movements (Fig. 5). In addition, we determined that this effect was due to changes in infraslow frequency correlations between pupil diameter and hemodynamics. This indicates that neurovascular coupling is heterogenous across cortex and brings to attention the fact that changes in CBV in prefrontal cortex is modulated by behavioral state.

While we saw a decrease in our hemodynamic signal in prefrontal cortex, we did not measure changes in oxygen concentration in these regions in response to an increase in orofacial movement which has been shown during voluntary movement periods to be modulated homogenously across cortex through an increase in respiratory rate (Zhang et al., 2019). Changes in arousal as measured by pupil diameter are also known to be an indicator of locus coeruleus activity which controls the release of norepinephrine (NE) through its expansive projections across the brain (Bekar, Wei, & Nedergaard, 2012; Reimer et al., 2016). Enhanced cortical NE is associated with a decrease in vessel diameter. Such vasocontraction could also drive the slight net negative correlation we observe between the two (Bekar et al., 2012; Turner, Gheres, & Drew, 2022).

In the context of rodent fMRI, most areas within the brain correlate negatively with changes in pupil diameter size at relatively short time lags, as do changes in calcium activity between 2-3 Hz (Pais-Roldán et al., 2020). However single trial analysis revealed that a fair bit of variability exists between this pupil-fMRI linkage highlighting the complex dynamic nature of arousal state on fMRI during resting state (Sobczak, Pais-Roldán, Takahashi, & Yu, 2021). In fMRI decreases in the BOLD signal are often interpreted as a decrease in neural activity in these areas with no consideration of behavioral state (Buckner, Andrews-Hanna, & Schacter, 2008; Fox et al., 2008). A mechanistic understanding of how fMRI recording of resting-state activity is influenced by arousal is critical both for the interpretation of future resting-state data and for understanding how cortical states influence behavior and functional network configurations (Drew et al., 2019). This will help us further understand to what degree resting-state spontaneous hemodynamic signal changes reflect local neural activity and non-neuronal responses (Winder et al., 2017). Future work should also carefully delineate the influence of cholinergic and adrenergic signals that control arousal on hemodynamic signaling. Studies to date suggest a significant spatial and temporal heterogeneity in this regard (DiNuzzo et al., 2019; Lohani et al., 2021; Schneider et al., 2016; Winder et al., 2017).

## Acknowledgments

We thank Dr. Thomas Knöpfel for sharing the VSFP 1.2 construct along with valuable advice on their use. We thank Dr. Hongkui Zeng for providing the initial VSFP mice for our colony. This work was supported by National Institute of Neurological Disorders and Stroke (UO1NS094302 and R01ns078095), National Institutes of Health BRAIN Initiative (R01 NS111470), and National Institute of Biomedical Imaging and Bioengineering (T32EB025816 and R01eb029857)

